# Epigenetic silencing and host genome evolution determine fate of viral insertions in *Acanthamoeba*

**DOI:** 10.1101/2024.10.31.621330

**Authors:** Cédric Blais, Morgan J. Colp, Luke A. Sarre, Alex de Mendoza, John M. Archibald

**Affiliations:** Department of Biochemistry & Molecular Biology, Dalhousie University, Halifax, Nova Scotia, Canada; Institute for Comparative Genomics, Dalhousie University, Canada; School of Biological and Behavioural Sciences, Queen Mary University of London, London, United Kingdom; Centre for Epigenetics, Queen Mary University of London, London, United Kingdom

## Abstract

*Acanthamoeba* is a cosmopolitan freshwater amoebae known for its association with Nucleocytoplasmic Large DNA Viruses (NCLDVs). Previous studies have shown that *Acanthamoeba* spp. undergo lateral gene transfer (LGT) with NCLDVs. Here we have leveraged chromosome-scale assemblies of two strains of *Acanthamoeba castellanii*, Neff and C3, to investigate the occurrence and genomic context of viral LGT in *Acanthamoeba*. We show that the viral ‘footprints’ in the C3 and Neff genomes are largely non-overlapping and that viral genes in Neff are found disproportionately in large sub-telomeric insertions. Multiple partial copies of these insertions are found throughout the Neff genome, but they are not expressed and do not encode functions for their own mobility. Viral regions are hyper-methylated and highly condensed, suggesting that the expression of recently acquired viral DNA is suppressed in heterochromatic regions. We propose a three-step model for the origin and fate of viral sequences in *Acanthamoeba*: (i) integration of DNA from giant viruses, (ii) epigenetic suppression of the viral DNAs, which allows them to persist in the genome, and (iii) deterioration of viral genomes by point mutation and intra- and inter-chromosomal recombination. Viral integrations in *Acanthamoeba* spp. are transient and may not have long-lasting effects on the fitness of the host amoeba. Our work strengthens a growing body of work showing widespread but transient integration of viral DNA in protists and extends the relevance of epigenetic silencing mechanisms to the evolution of *Amoebozoa*. We highlight the importance of host genome dynamics for understanding the evolution of endogenized viral elements.

## Introduction

The acquisition and spread of mobile genetic elements are well-known drivers of genome evolution in eukaryotes. Large stretches of the genomes of plants, animals and fungi consist of so-called ‘junk’ DNA, which for the most part does not encode proteins used by the host cell and whose sequence evolves under relaxed selective constraints^1^. The role this junk DNA plays in the organism is hotly debated^2,3,4^. It is often considered a by-product of the spread of selfish DNA, such as transposable elements and retroviruses, which replicate to their own advantage without benefitting the host genome. The factors that lead to the acquisition and retention or loss of virus-derived DNA remain poorly understood. Recent developments in comparative genomics show that we have much to learn from the realm of microbial eukaryotes, in which new classes of mobile and viral elements have been discovered. These elements form symbiotic and parasitic relationships with distant lineages of eukaryotes, connecting them into a complex genomic ecosystem which provides numerous avenues for lateral gene transfer (LGT) within and between branches of the eukaryotic tree of life^5,6,7^.

Members of the virus kingdom Bamfordvirae are emerging as a major source of foreign DNA in eukaryotic genomes^8^. The Bamfordvirae are incredibly diverse dsDNA viruses characterized by their use of double jelly roll major capsid proteins^9^. They notably include ‘giant’ viruses, also known as Nucleocytoplasmic Large DNA Viruses (NCLDVs). Comparative genomics suggests that the group has evolved viral gigantism multiple times^10^, with the genomes of some NCLDVs being as large as 2.6 megabase-pairs (Mbp)^11^. NCLDVs have been shown to both donate genes and receive genes from eukaryotic genomes, in which they reside in various states of degradation^12,13^. While most cultured representatives of NCLDVs are lytic, this is possibly because they are typically isolated by inoculating protist cultures and monitoring them for lysis^14^. Interestingly, a 300 Kilobase-pair (Kbp) endogenous mirusvirus – a chimeric relative of NCLDVs^15^ – was recently found integrated in the genome of the thraustochytrid protist *Aurantiochytrium limacinum* (the organism also harbours a similarly-sized, evolutionarily distinct episomal version of the same virus)^16^. Similarly, a 617 Kbp integrated NCLDV was discovered in *Chlamydomonas reinhardtii* which could be induced to produce virions^17^, raising the possibility of lysogenic cycles being widespread among NCLDVs. In addition to these endogenized giants, an extensive network of modular, interconnected and highly diverse mobile elements has been shown to integrate into protist genomes^18^. It includes virophages, superparasitic viruses who target other viruses and may form mutualistic relationships protecting the cells in which they are integrated, and polinton-like viruses, which may alternate between mobile-element-like lifestyles and viral lifestyles^7^, and may have evolved their own form of superparasitism^19^.

The long-term effect of DNA viruses on the genome biology and evolution of eukaryotes remains unclear and is likely highly variable across the eukaryotic tree. Viral integrations do not always necessarily translate into lasting contributions to host genome evolution. Viral insertions vary greatly from one genome to the next^20^ and it can be difficult to distinguish ‘live’ endogenized viruses which retain virulence from permanently integrated sequences co-evolving with the genome. While some protein-coding genes in eukaryotes appear to be the product of ancient LGT from viruses^21^, for the most part viral contributions to eukaryotic genomes appear to be recent and transient^22,20^.

The genus of single-celled amoebae *Acanthamoeba* has been particularly fruitful for the study of the interplay of eukaryotes and Bamfordvirae. *Acanthamoeba* are hosts to a broad range of intracellular bacteria and viruses and have been referred to as melting pots of LGT^23^. The first recognized ‘giant’ virus, Mimivirus, was isolated from *Acanthamoeba polyphaga* in 2003^24^, and *Acanthamoeba* has since proven to be permissive to an extremely broad range of NCLDVs^25^. This permissiveness has led it to be used to discover and isolate many new lineages of viruses^26^, including virophages, which were found to co-occur with giant viruses in *Acanthamoeba*^27^. The *Acanthamoeba* genome bears the mark of these associations – hundreds of genes derived from viruses have been identified in several species of *Acanthamoeba*. For example, Maumus and Blanc (2016) identified several large, poorly expressed clusters of viral sequences in the original reference genome assembly of *A. castellanii*^28^ and provided evidence for collinearity between these viral integrations and some giant viral genomes^29^. However, the limited number of then available Bamfordvirae genomes restricted the search for viral sequences, and the fragmented nature of the assembly prevented examination of the genomic context in which these transfers occurred. This made it difficult to draw inferences about the evolutionary trajectory of giant viral integrants in *A. castellanii* strains and related species.

We have performed a genome-wide investigation of viral DNA in two recently published chromosome-scale assemblies for *Acanthamoeba castellanii* strains Neff and C3^30^. The high quality of these assemblies provides a unique opportunity to explore the genomic context of viral integrants in these two divergent strains, thereby allowing us to uncover factors that shape their acquisition and fate in a nuclear context. Our results suggest that epigenetic suppression of viral DNA expression is an important factor limiting deleterious impact of newly integrated viral DNAs, as is recombination-based disruption of intact viral genomes.

## Materials and Methods

### Detection of viral sequences

Genome assemblies and gene models for *Acanthamoeba* strains C3 and Neff used in this study were from Matthey-Doret, Colp et al.^30^. ORFs 150 nucleotides and longer were extracted from intergenic regions using ORFm^31^ and were not required to have a start codon. Predicted proteins and intergenic ORFs for Neff and C3 were BLASTed against a local copy of the NCBI nr database (downloaded May 2024) using Diamond BLAST^32^ with an e-value cutoff of 0.001. All hits to the *Acanthamoeba* genus were filtered out to avoid missing recent LGTs. Protein sequences with a best hit to viruses, or which had viruses as half of their top ten hits, were retained as viral candidates and mapped to the Neff and C3 assemblies. ViralRecall was also used to cross-reference our viral candidates^33^.

### Analysis of genome conservation between Neff and C3

A whole genome alignment of the Neff and C3 assemblies was performed using MUMMER4^34^ with an extend value of 1,000 bp to avoid losing alignments due to short gaps. Candidate viral proteins were identified as being conserved between the two genomes if they mapped wholly or partially to regions that could be confidently aligned between the Neff and C3 genomes. They were identified as not conserved if they completely overlapped with a gap in the alignment. The show-diff function was used to identify differences between the Neff and C3 assemblies and identify possible duplications and translocations. Due to the prevalence of small (a few hundred to a few thousand base pairs) translocations, possible translocations were split between small translocations below 20 kbp, which are not indicative of large-scale chromosomal reorganization, and larger translocation events. The boundaries of viral integrations were difficult to identify, as gaps in the Neff-C3 alignment corresponding to viral ORFs were often interrupted by duplications and/or translocations from elsewhere in the genome. A custom script was written which concatenated together contiguous gaps, translocations and duplications. The coordinates of these concatenated divergent regions were used as a proxy for the beginning and end of viral integrations.

### Comparison of viral insertions within the Neff and C3 genomes

All concatenated divergent nucleotide regions coding for putative viral proteins were BLASTed^35^ against one another, using the DUST program to filter out hits to low-complexity sequences. Self-hits were removed and the interconnections between concatenated divergent regions were visualized as a sequence similarity network using Cytoscape^36^. A more detailed alignment was then performed for selected viral insertions using Sibelia^37^ and Circos^38^.

### Functional annotation and detection of mobile elements

Predicted functions were assigned to all Neff and C3 predicted proteins and intergenic ORFs using an InterProScan^39^ search with an e-value cutoff of 0.001. To further analyse the completeness of lineage specific viral insertions, the Neff and C3 genomes were searched for viral hallmark genes using ViralRecall. Hits with an e-value below 1 × 10^−5^ and which were found in a region not aligning to the other genome were retained. The InterProScan annotations for viral candidates were manually examined for functions indicative of mobility. Mobile elements and simple repeats in Neff and C3 were predicted using RepeatMasker^40^. EDTA (Extensive de novo TE Annotator)^41^ was also used to predict the location of mobile elements as well as terminal inverted repeats.

### Expression and repression of viral sequences

RNA-Seq reads from strain C3 (uninfected with *Legionella*) from Matthey-Doret, Colp et al.^30^ were combined into one file and aligned to the C3 genome assembly using HISAT2^42^. Neff RNA-Seq reads were similarly aligned to the Neff assembly. Transcript Per Million (TPM) values were calculated with Geneious Prime 2023.04 (https://www.geneious.com) for all predicted proteins and viral intergenic ORFs. Hi-C data generated by Matthey-Doret, Colp et al.^30^ was recovered from https://zenodo.org/records/6800059 (31/10/2024) to identify chromosome territories and patterns of DNA packaging associated with viral insertions. 5-Methylcytosine base modifications in Neff and C3 were predicted using the nanopolish call-methylation package^43^ and the raw fast5 reads underpinning each assembly. Neff nanopore fast5 files were generated in the Archibald Lab and the nanopore fast5 files for C3 were provided by Cyril Matthey-Doret, stemming from Matthey-Doret, Colp et al.^30^.

### Controlling for mis-assemblies and allele-specific variants

Mis-assemblies can be a major confounding factor when analysing spatial distribution patterns of viral DNAs in nuclear genomes. To control for this fact, nanopore long reads from both the Neff and C3 assemblies were mapped against the final assemblies using minimap2, and Hi-C contact data generated by Matthey-Doret, Colp et al.^30^ were considered for all chromosome-scale scaffolds. We manually inspected both the long read alignment and the Hi-C plots to identify possible mis-assemblies. Genomic regions were considered to be possible mis-assemblies if a sequence’s location lacked both long read mapping and Hi-C support. Spatial patterns in the distribution of viral genes and translocations were filtered to exclude sequences falling within these compromised regions and reanalyzed to show that observed biases still held. In several cases, mis-assemblies appeared to be a product of allele-specific variants not being properly reflected in the assembly and in some cases artefactually grafted onto the ends of scaffolds. The true location of these variants could be resolved using Hi-C data and by mapping reads to their true location.

## Results

### The genomes *of Acanthamoeba castellanii* strains Neff and C3 both harbour hundreds of sequences from giant viruses

We identified 750 viral LGT candidates in Neff and 642 in C3 (Fig. 1), a substantial increase from the 267 reported by Maumus and Blanc (2016)^29^ in Neff and the 115 found by Chelkha et al. (2018) in *A. polyphaga*^44^. This increase is in large part likely due to improvements in the sampling of giant viruses, as genomic data from new isolates can dramatically reshape the taxonomic landscape of viral insertions. For example, the most common viral best hit using the Neff genome as query was to Medusavirus, a lineage discovered in 2019. Many of the viral LGT candidates in Neff and C3 are found in viral ‘islands’ where they co-occur with orphan genes of similar size and with few introns. Since ORFans are very common in NCLDV genomes^11^, we expect that viral homologues will be identified for many of these endogenized sequences as sampling of giant virus genomes continues to improve. Our expanded search criteria also explain some of the increase in identified viral sequences. Our search for viral sequences in intergenic regions included all ORFs 50 amino acids or longer, regardless of whether a start codon could be found. As a result, some groups of viral ORFs are predicted to be pseudogenes, with each ORF mapping to a different segment of the same viral protein. This degraded state suggests that some viral sequences are evolving under relaxed selective constraints.

**Fig. 1.**
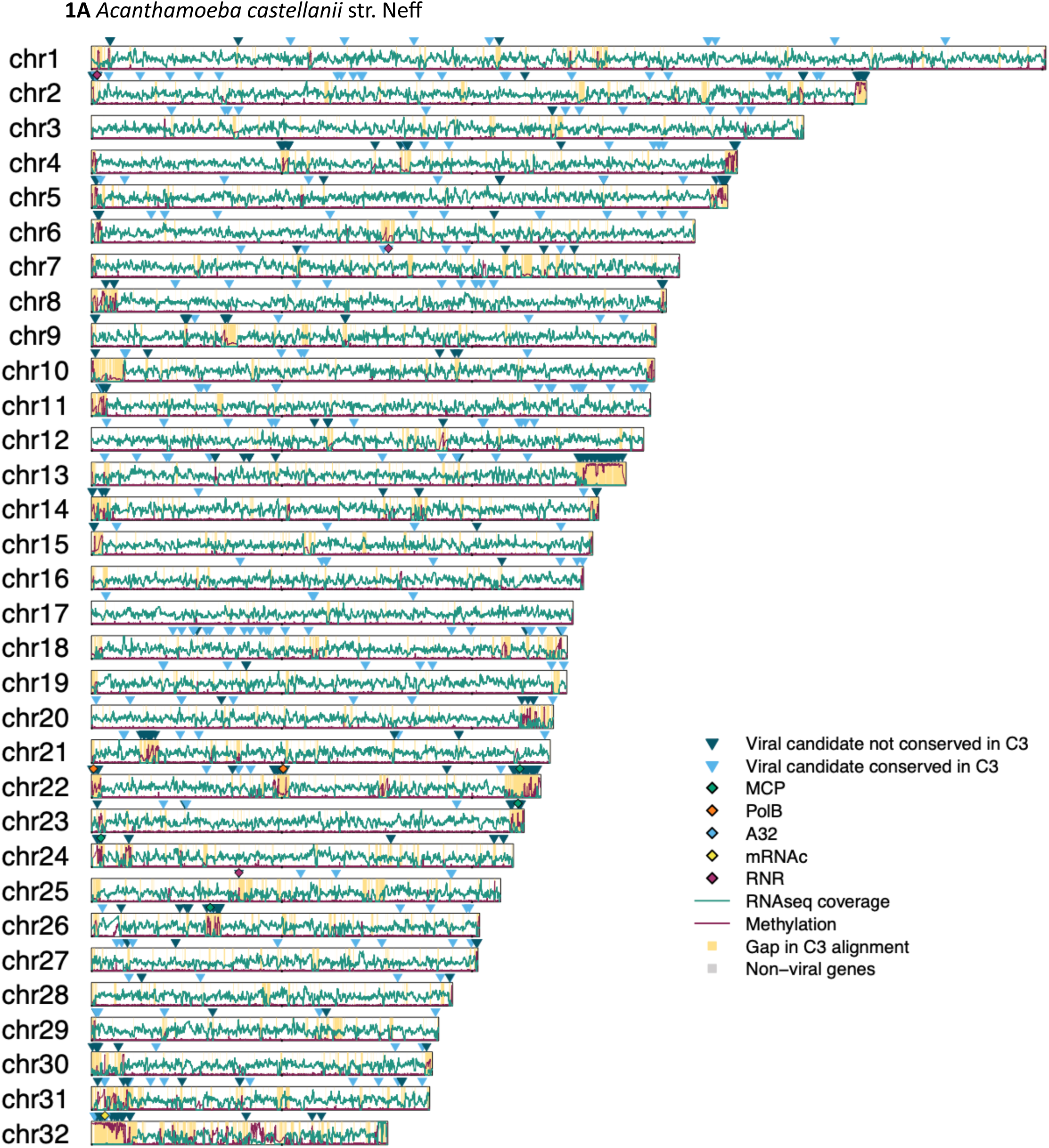
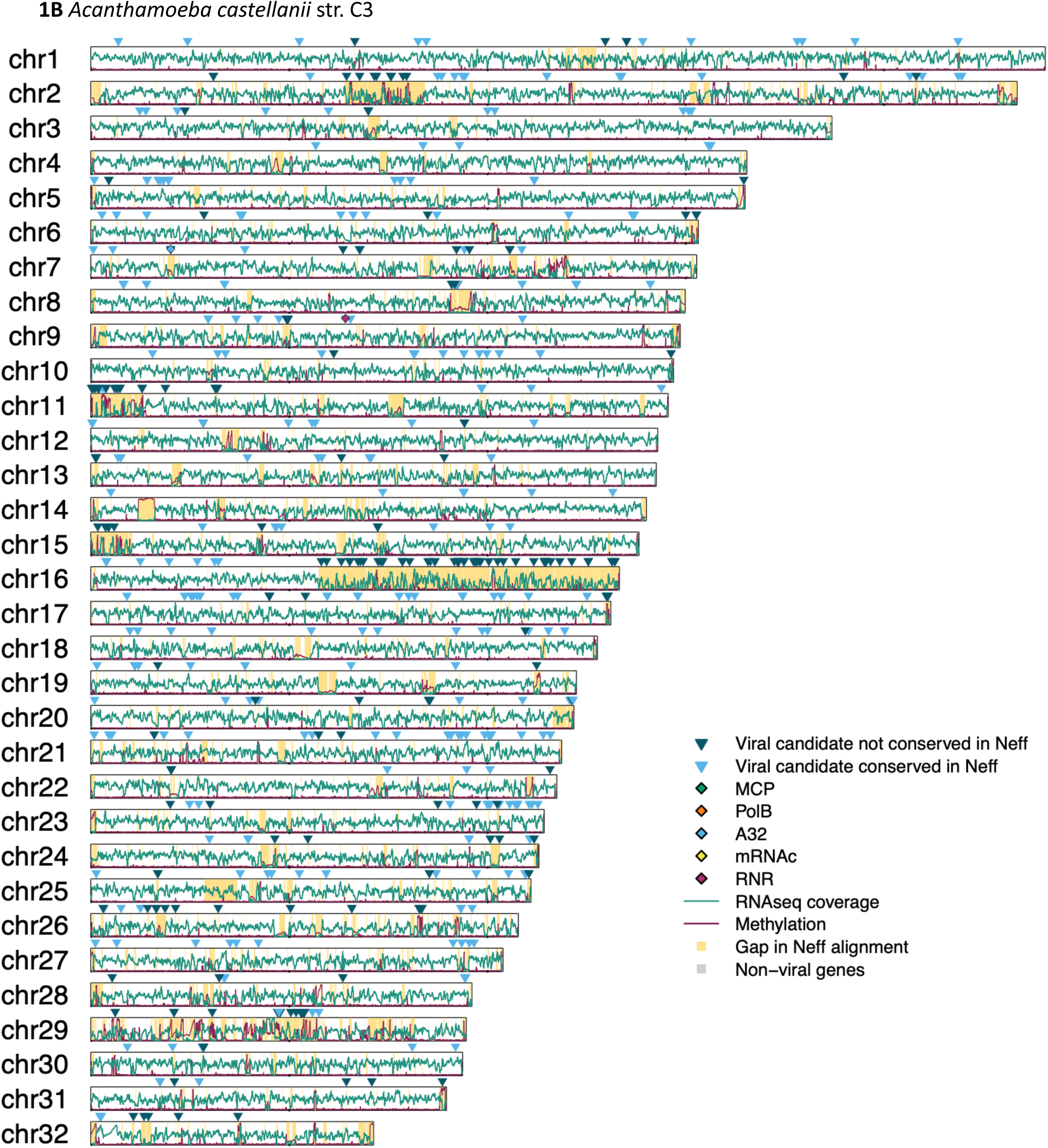
Location of viral genes, methylation levels, expression levels, and gaps in alignment across the first 32 chromosomes of *Acanthamoeba castellanii* strains Neff (A) and C3 (B).

### Viral proteins are clustered in poorly conserved insertions, and near sub-telomeric regions in the Neff genome

Putative virus-derived genes in Neff tend to cluster together, forming viral regions usually not conserved in C3. This suggests that they are the product of strain-specific insertions of viral DNA chunks carrying multiple genes. Similar insertions are also found across C3, though they tend to be smaller and encode fewer viral ORFs (Fig. 1). The observed differences in level of clustering are likely partly influenced by the higher number of lineage-specific insertions in Neff. 460 of the 750 viral candidate proteins in Neff do not have an obvious counterpart in C3, whereas 301 of 642 viral candidate proteins in C3 were not conserved in Neff. Conversely therefore, more viral candidate proteins in C3 were conserved in Neff than vice-versa. The discrepancy in the number of conserved regions between Neff and C3 is in part due to the use of nucleotide level alignment of genomic regions as a proxy for inter-strain conservation. For example, a region encoding a putative viral ORF in Neff might align to a region in C3 of likely viral origin, but where no viral ORF was detected, possibly due to partial deletion of the viral sequence. Likewise, strain-specific duplication of viral regions would cause two regions of one genome to map to only one in the other. These patterns could also be explained in part by secondary loss in Neff and/or C3, but in any case, the lack of conservation of viral DNA insertions between the two stains suggests that they are limited in their taxonomic range.

Only one cluster of viral ORFs is conserved between Neff and C3. Conserved viral candidates in Neff and C3 tend to be isolated from other viral candidates in terms of their genomic locations and are usually larger genes with more spliceosomal introns. This is consistent with the hypothesis that most evolutionarily conserved ‘viral’ candidates in *A. castellanii* are in fact eukaryotic genes that were acquired by viruses and not vice versa. A good number of them are annotated as serine/threonine kinases and are part of a large family of kinases with a unique domain architecture that is shared between *Acanthamoeba* and giant viruses^28,29^. Phylogenetic trees of these proteins (data not shown) suggest that they have been transferred from nuclear genomes to viruses and back again multiple times, and that their prevalence in the Neff and C3 genomes is more a product of gene duplication common to kinase families rather than recent re-acquisitions.

Insertions in Neff are disproportionately found near the ends of chromosome-scale scaffolds and close to both canonical (TTAGGG) and degenerate telomeric repeats. Interestingly, translocations relative to C3 are also disproportionately found in sub-telomeric regions of Neff chromosomes (Fig 2). While sub-telomeric regions in Neff were found to contain numerous mis-assemblies, this spatial bias was observed even after manually curating the Neff scaffolds to remove assembly artefacts. It is unclear whether a similar bias is found in C3, which has fewer large insertions on chromosome-scale scaffolds, making overall trends difficult to assess. This difference is likely in part due to the smaller number of lineage specific insertions in C3, and to assembly artifacts. Indeed, viral candidates in C3 are disproportionately likely to be found on small scaffolds whose chromosomal location is unclear, at least one of which contains degenerate telomeric repeats. Notably, three of the four largest insertions on chromosome scale scaffolds are flush with the end of a scaffold (see chr11, chr15, and chr16 in Fig. 1B). It is thus plausible that C3 harbours a similar sub-telomeric bias, but that a paucity of insertions and assembly limitations have made it less detectable.

**Fig. 2.**
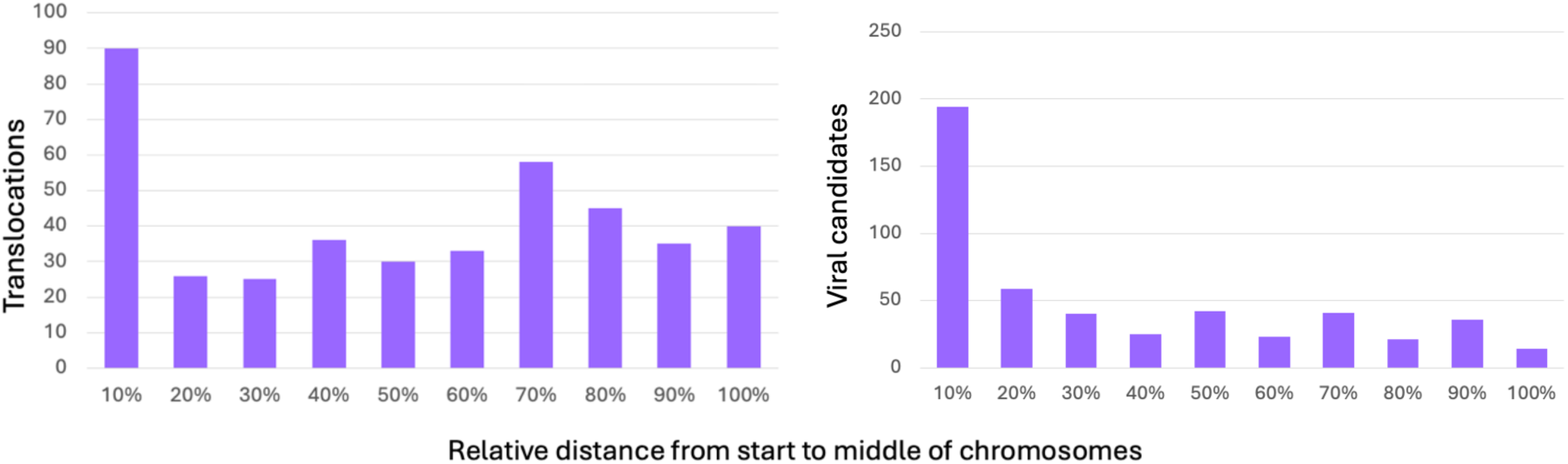
Relative distance of translocations (A) and viral candidates (B) from the start of chromosomes, 10% corresponding to being within 10% of half a chromosome’s length of the end, and 100% being in the middle of the chromosome.

### The C3 genome harbours a large chimeric region which may have triggered large-scale chromosomal recombination

A striking feature of the C3 viral footprint is the large-scale chimerism of chromosome 16. Two thirds of the chromosome’s 1.3 Mbp sequence, starting at 575 Kbp, is not conserved in Neff (Fig. 1B, chr16). The anomalous, non-conserved half of the chromosome is gene poor and enriched for viral candidates, although these are not as densely clustered as viral homologs in Neff and are interspersed with sequences of apparent eukaryotic origin. Transcription levels are lower and more heterogenous than in the more typical eukaryotic half. Interestingly, a search for viral hallmark genes in this stretch of C3 chromosome 16 using ViralRecall yielded no results. The GC content is unchanged compared to the rest of the genome however, and the junction between the ‘normal’ and ‘chimeric’ halves of the chromosome is well supported by long-read data, although multiple breaks in the read coverage can be found starting at ∼830 Kbp. The physical proximity of regions separated by breaks in the long-read coverage is supported by Hi-C data, suggesting that they are part of the same molecule and not the product of an assembly artefact. There is some evidence that this naturally chimeric region has been the catalyst for large-scale chromosomal reorganization, as the non-chimeric half of chromosome 16 in C3 maps to chromosome 2 in Neff. In contrast, chromosome 2 in C3 maps both to the other half of chromosome 2 in Neff as well as Neff chromosome 25, although curiously this chimerism with chromosome 25 is supported only by Hi-C and not read data. Hi-C data further suggest that the chimeric portion of C3 chromosome 16 has some connection to a region in chromosome 2 that is likewise supported by Hi-C but not long-read data. While unresolved mis-assemblies prevent us from drawing a definitive conclusion about the origin of the chimeric chromosome 16 in strain C3, on balance the evidence suggests that it is genuine. *Acanthamoeba* is both highly polyploid and aneuploid^45^ and so the association between a massive chimeric region on chromosome 16 and the only large-scale difference between the Neff and C3 assemblies is likely due to the real underlying biology of *Acanthamoeba*.

### Neff and C3 viral proteins have conventional viral functions but insertions in both genomes lack a full suite of viral hallmark genes

Several viral ‘hallmark’ genes, which are highly conserved in NCLDVs and involved in key viral processes, were identified in both genomes. Those include several major capsid proteins, viral polymerase B, and A32 packaging ATPase, among others. Notably, no viral insertion in either genome sported an obvious full suite of viral hallmark genes (Fig. 1), and some hallmark genes (the Viral Late Transcription Factor 3 (VLTF3) and Superfamily II Helicase (SF II)) were missing from all insertions in both Neff and C3. An InterPro search failed to recover a potential function for 380 of the 750 putative viral candidate proteins in Neff. Of those that remained, the most common functional annotation was that of serine/threonine kinase, which is part of a large family of kinases shared with viruses, discussed above. Furthermore, no mobility-associated functions were identified in the viral proteins encoded by either genome, which suggests that these insertions are not capable of mobilising by themselves in the *Acanthamoeba* genome.

### Viral insertions are dynamic, often chimeric, and sometimes allele-specific

The structure of viral insertions in the Neff genome is complex. Viral insertions are interspersed with nucleotide sequences conserved in C3 and duplicated or translocated from elsewhere in the genome, as well as sequences not conserved in C3 but with copies in other locations in the Neff assembly. These relocated sequences can have wildly different expression levels and GC contents compared to the insertions in which they are embedded. Post-insertion recombination dynamics likely scramble the boundaries of viral regions and can make it difficult to tell where inserted viral DNA ends and the recipient eukaryotic genome begins.

The difficulty of delineating viral insertions is compounded by the fact that they may themselves be dynamic to some degree. An analysis of nucleotide sequence alignments of strain-specific sequences encoding putative viral proteins in Neff shows that multiple viral insertions in Neff share regions with nucleotide identity around 85-95%, comparable to the 85% average sequence identity between Neff and C3. These duplicated regions range from a few hundred to several thousand nucleotides, and typically correspond to a subset of a larger, non-duplicated viral region. The presence of these duplicated viral regions in Neff cannot obviously be attributed to any intrinsic mobility of viral insertions. As discussed above, no mobility-associated functions were found in the InterPro search for viral candidates, and the possible borders of viral insertions do not obviously correspond to terminal inverted repeats. This, along with viral insertions’ inconsistent size, gene content, and apparent affiliation with giant viruses speaks strongly against them being transposable elements or polinton-like viruses and virophages.

Instead, duplicated viral sequences could be products of multiple insertions from giant viruses and/or the duplication and dispersal of viral DNA following insertion in the genome. It is difficult to decisively rule for one against the other, however the dynamic genomic context of viral insertions lends credence to the notion that at least some duplicated viral regions are the product of post-insertion duplication. Non-viral Neff sequences that are conserved in C3 but which have since duplicated or translocated are often found near or even within viral insertions. Likewise, several viral insertions in Neff share long stretches of DNA with no apparent viral signature, which may correspond to transposable elements. At least one viral region contains an insertion from an LTR retrotransposon (Fig. 3). Together these observations suggest that the apparent mobility of viral insertions in Neff is due to processes intrinsic to the host nuclear genome, such as mobile elements and translocation. This inference is further supported by the observation that translocations in Neff are disproportionately found in sub-telomeric regions (Fig. 2). The sub-telomeric bias of viral insertions reflects broader trends in the genome’s evolution.

**Fig. 3.**
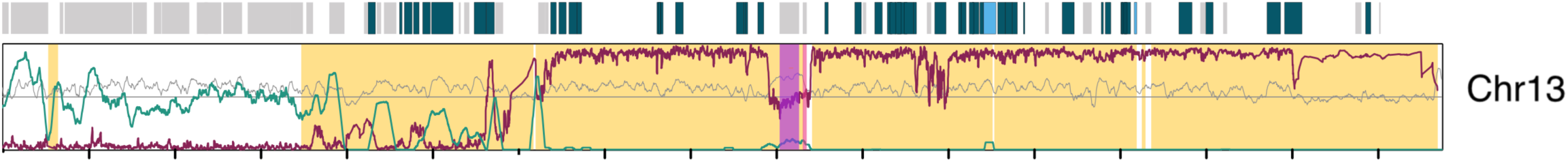
Detailed examination of the chimerism and complexity of a large viral insertion on Chromosome 13 of *Acanthamoeba castellanii* str. Neff. Dark blue-green boxes on top indicate putative viral genes not conserved in C3. Light blue boxes indicate putative viral genes with some nucleotide alignment to C3. Gray boxes indicate genes not flagged as viral. A reverse transcriptase-encoding ORF and integrase-encoding ORF with counterparts elsewhere in the genome are highlighted purple and pink, respectively. Methylation level is shown in red, expression in green. Regions not conserved in *A. castellanii* strain C3 are shown in yellow. GC content is shown in grey.

In addition to this duplication process, recombination between viral insertions with distinct origins can also be inferred from alignment data. Sudden shifts in GC content are commonplace in viral insertions in Neff. For example, a large sub-telomeric viral region on Neff chromosome 5 is split into at least two regions with differing GC contents, one of which aligns to a viral insertion on chromosome 21, and the other aligning to a viral region on chromosome 2. While duplication or multiple integrations of an already heterogeneous viral insertion followed by differential loss could explain this pattern, post-insertion chimerism is consistent with the observation of transposable elements, duplications, and translocations in the vicinity of viral regions.

Finally, some viral insertions also appear allele-specific in both Neff and C3, with varying degrees of post-insertion duplication and recombination events between alleles. Allele specificity is inferred when an insertion is well supported by some, but not all, reads mapping to the chromosome, indicating the presence of multiple variants of the same position in the genome. For example, a large insertion at the end of Neff chromosome 22 corresponds to a sudden drop in depth of read coverage corresponding to a split between reads that either map or do not to the reference assembly. Many of the non-reference reads end in telomeric repeats, indicating that they correspond to poylmorphic sub-telomeric regions lacking a viral sequence.

A large viral region spanning ∼100 Kbp at the end of Neff chromosome 13 captures these multiple layers of complexity. The region starts with a ∼12 Kbp segment coding for multiple viral proteins, which is an almost complete copy of a putative sub-telomeric insertion on chromosome 6. The duplicated region is less methylated and more expressed than the main body of the viral region. The latter contains an insertion by an LTR retrotransposon with a distinct GC content (68%, compared to 57% in neighbouring sequences). Other copies of this transposable element are found in the genome, which likely integrated in the viral DNA following its insertion in Neff. Finally, the last segment of this large viral region has a distinctively lower GC content (∼50% instead of ∼57%) and is also supported by only a subset of the reads mapping to that location – indicating that this additional sequence was likely grafted onto some, but not all, alleles of this viral region (Fig. 3).

### Viral insertions in the Neff and C3 genomes are transcriptionally suppressed by methylation and heterochromatin

Viral insertions in both Neff and C3 are transcriptionally suppressed. By analyzing 5-methylcytosine methylation signal from nanopore data (from Matthey-Doret, Colp et al.^30^), we found that non-conserved regions in general account for most spikes in 5-methylcytosine density in both genomes, and viral insertions in particular show the highest methylation levels (Fig. 1). Furthermore, the Hi-C contact maps of the Neff genome show that some of the largest viral insertions correspond to densely packaged chromosomal territories (Fig. 4), unlike regions of clear eukaryotic origin. Similar, though less distinct, territories can be observed in C3. The most striking viral chromosomal domains correspond to the two largest viral insertions in Neff, at the end of chromosome 13 and start of chromosome 32. Chromosome 32 is believed to be the home of the Neff 18S and 28S ribosomal RNA (rRNA) genes, and inter-chromosomal contact maps show that it is enriched for physical contact with most other chromosomes^30^. There is extensive contact between the viral insertions on chromosome 13 and 32 on a scale not observed elsewhere in the genome (Fig. 4). The distinct repression status of these two inserts is further supported by their high methylation levels. Viral inserts in both Neff and C3 typically show high yet uneven levels of methylation, with abrupt changes in the methylation level of integrated sequences. Methylation levels in the chromosome 13 and 32 viral insertions are comparatively consistently high over long stretches (Fig. 1). It is likely that the two viral regions are packaged together as part of a single heterochromatic region.

**Fig. 4.**
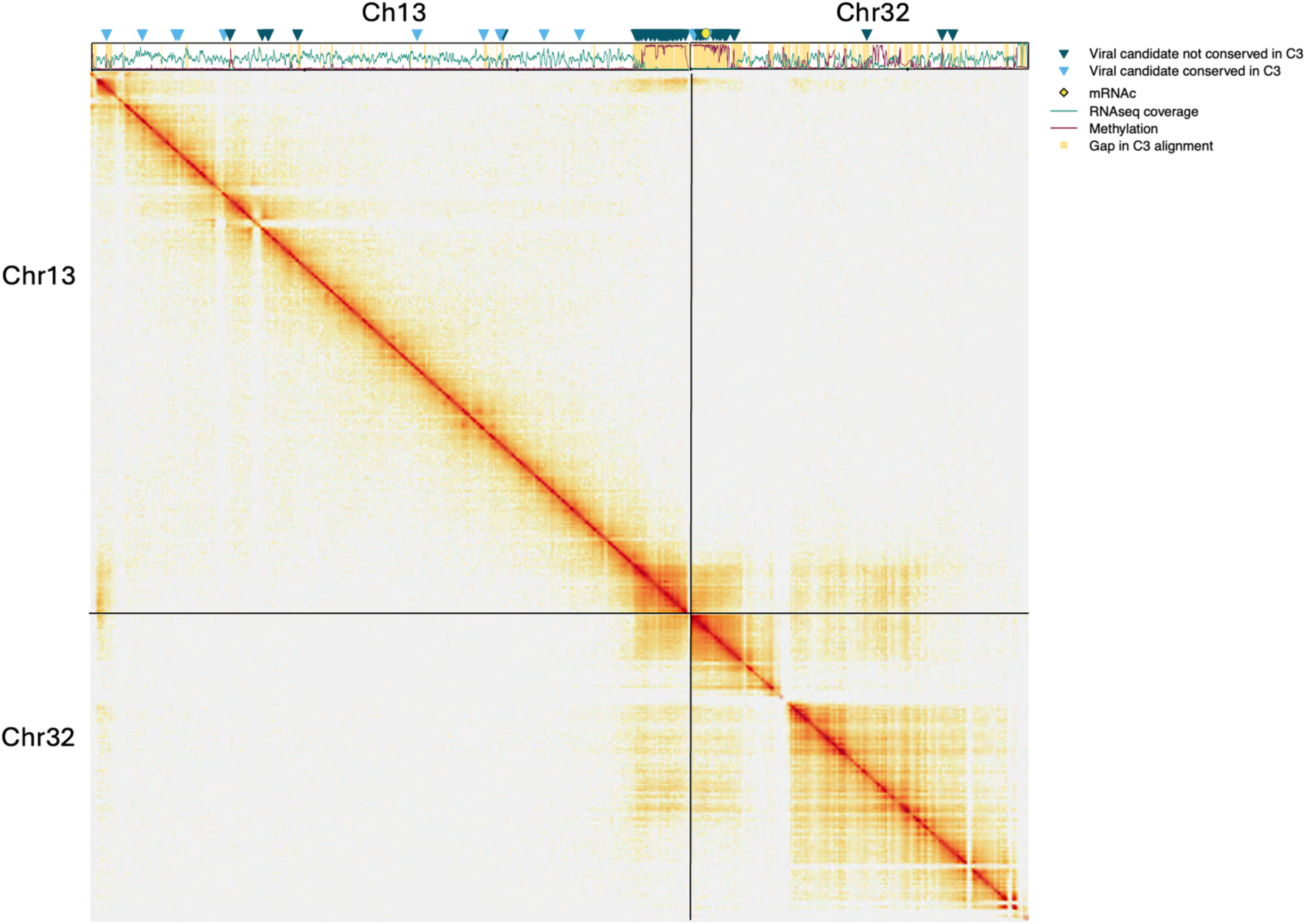
Hi-C data plot showing intra and inter-chromosomal contacts between *Acanthamoeba castellanii* Neff chromosomes 13 and 32. The viral candidate mapping is shown at the top.

## Discussion

### Viral insertions in Neff and C3 are transient and likely non-functional

We have performed a genome-scale analysis of the origin and fate of viral DNAs inserted into the genomes of two strains of *Acanthamoeba castellanii*, an organism that has featured prominently in the field of giant viral research. Our study contributes to a growing body of literature showing that NCLDVs have an extensive but often transitory footprint in protist genomes^20,22^. Both fragments and full NCLDV genomes have been found integrated in the nuclear genomes of a wide range of unicellular eukaryotes^13,44,46^. Despite some ancient and long-lasting contributions of giant viruses to the eukaryotic proteome^21^, observed integrations tend to be lineage specific and vary greatly between lineages and even between closely related species^20^. Our findings are consistent with this pattern. Despite their substantial genetic differences, Neff and C3 are generally considered to be strains of the same species; they share 97% 18S rDNA identity and ∼85% of their orthogroups^30^. The average sequence identity of the whole genome alignment is ∼85%. Only 290 of 750 viral candidates in Neff and 341 of 642 of viral candidates in C3 are conserved between the two, a large proportion of which are likely *bona fide* eukaryotic genes with viral homologues due to eukaryote-to-virus transfer. Most viral regions appear to have been acquired after the divergence of the two strains, although we cannot rule out ancestral acquisition and differential loss, the signal for which would be erased due to rapid sequence divergence. The pseudogenization of several viral genes and the observation of allele specific recombination events in viral insertions suggests they are evolving under relaxed selective constraints. Furthermore, the active suppression of these viral elements by the cell through methylation and chromatin condensation further buttresses the case that viral integrants are not useful to the cell.

### Viral DNA acquisition in *Acanthamoeba*: endogenization, suppression, degradation

Whether the widespread presence of viral fragments in protist genomes is a product of accidental or deliberate insertions that are part of the viruses’ life cycle cannot be answered decisively without knowing the identity of the viral donors. As it stands, most viral candidate proteins share low sequence identity to their best viral hit, indicating that closer relatives – whose lifestyles are unknown – may yet be found. We propose a three-step model to explain the widespread discovery of chunks of incomplete viral sequence in the genomes of *Acanthamoeba* and other lineages: (i) integration of viral DNA into the genome, (ii) epigenetic suppression of endogenized viral DNA, and (iii) disruption of the viral DNA by mutational processes.

#### Step 1: Nuclear association provides a pathway for viral DNA integration into the host genome

Fragmented viral insertions are observed in a wide range of eukaryotic genomes. Whether these insertions are a product of ‘accidental’ integration during failed infection and/or occur due to endogenization of full viral genomes as part of a latent infection cycle is debatable. In *Acanthamoeba castellanii* at least, this question cannot be resolved without knowing the identity of the viral donors. Multiple capsid proteins encoded by viral insertions in Neff have a best hit to Medusavirus, with amino acid identities as high as 75%. Most hits are far weaker, however. While Medusavirus is the most common best hit for Neff viral proteins, Pandoravirus is also well represented, and top hits to several other viral lineages can also be found. Interestingly, in strain C3, Pandoravirus is the most common best hit, with Medusavirus being second.

NCLDV infection is a highly disruptive process. In the case of Mimivirus, the first discovered giant virus^24^, which is known to infect *Acanthamoeba*, the cell nucleus shrinks dramatically during infection even though viral replication occurs entirely in a cytoplasmic viral factory^47^. Pandoravirus, on the other hand, requires entry of the viral genome into the nucleus for replication, followed by the formation of a viral factory in the cytoplasm and degeneration of the nuclear membrane^47^. Medusavirus replicates inside of the host nucleus and appears to leave the nucleus morphologically intact until later in the infection process, which often results in encystment of the cell^48^. Given these observations, close contact with and disruption of the *Acanthamoeba* nucleus may result in accidental integration of viral DNA fragments into the nuclear genome. Yet to be discovered non-lytic relatives of Medusavirus and Pandoravirus could also have colonized the *Acanthamoeba* genome. While the majority of known NCLDVs are lytic, current NCLDV sampling methods rely on the observation of cell lysis^14^ and as such bias discovery towards the most lethal viruses. With the increasing number of examples of fully endogenized, activatable NCLDV and NCLDV-like genomes in protists and algae ^16,17^, it is possible that the true missing, non-lytic viral donors in *Acanthamoeba* exist but have not yet been identified, despite the widespread use of this amoeba in NCLDV research. Either way, the identification of endogenized viruses also related to Medusavirus in the distantly related opisthokont *Amoebidium*^22^ supports the case that the Mamonoviridae lineage or its close allies are prolific viral DNA donors.

#### Step 2: Suppression allows viral DNAs to remain in the genome over evolutionary timescales

The pervasiveness of viral methylation in *Acanthamoeba* raises questions about the nature of host-virus interactions. Most methylation in Acanthamoeba targets NCLDV sequences. This could suggest that NCLDV integration represents a common enough threat to warrant a dedicated cellular response, buttressing the case that integrations are degenerate products of endogenized, latent viruses. This is not necessarily the case, however, as some recent transposable elements in the Neff genome are also hypermethylated. Hypermethylation of viral insertions may thus be a byproduct of more generic self/non-self defence systems targeting any foreign DNA in *Acanthamoeba*, which would affect even rare accidental integration events.

Whether viral DNA integration in *Acanthamoeba* is accidental or not, its silencing by 5-methylcytosine base modification likely facilitates the persistence of viral insertions over evolutionary timeframes. If viral DNA enters the genome because of an endogenous ‘lifestyle’, silencing it will reduce the likelihood that the virus kills the cell. In this way suppression reduces the fitness cost of the inserted virus while also allowing it to persist in the genome over multiple generations. These fitness benefits also likely apply in cases of accidental integration. Viral infection results in substantial host cell reorganization; Pandoravirus recycles the nuclear membrane for virion formation^49^, diverse NCLDVs encode a wide range of metabolic genes to reprogram the infected host cell^50^, and even cellular locomotion can be altered, as the Mimivirus relative Tupanvirus induces infected amoebae to seek out and aggregate with non-infected cells^51^. The expression of even an incomplete viral proteome could impact host fitness, a fitness cost which methylation reduces regardless of whether it evolved specifically to target viruses. In this scenario, the widespread distribution of virus insertions could be a mere byproduct of the reduced fitness cost of rare insertions, thanks to non-specific methylation of foreign DNA.

Our results are broadly consistent with work on the opisthokont *Amoebidium*, where large hypermethylated islands coincided with partial NCLDV integrations. Moreover, other unicellular holozoans showed a correlation between 5-methylcytosine modifications and the presence of giant virus endogenizations^22^. Similarly, giant virus endogenizations were found to be methylated in the moss *Physcomitrium patens*^52^. 5-methylcytosine appears to enable the co-evolution of selfish DNA with the genome by reducing the fitness cost of genetic parasites. Our account of hypermethylated giant virus insertions in *Acanthamoeba* strengthens the applicability of this model to *Amoebozoa* and shows that methylation enables similar patterns of integrations in distantly related organisms.

#### Step 3: Degradation neutralizes the virus and produces observed patterns of scattered viral insertions

Host-associated molecular silencing mechanisms that limit translocation and mobile element spread facilitate the final step in the inserted DNA’s evolutionary trajectory. Once silenced, the viral DNAs are at the mercy of mutational processes which will mutate and re-arrange them, inactivating any complete viral genomes and degrading individual protein-coding sequences before eventually deleting them. These processes are on display in the *A. castellanii* Neff genome, where partial copies of viral insertions are found scattered throughout the genome. Examples of deletions and recombination can readily be observed across duplicated viral regions, as can be colonization of viral insertions by sequences and transposable elements from the host genome. Once broken down as such, the virus seems far less likely to have a fitness effect on the cell, which nonetheless continues to repress it. The sub-telomeric bias in viral integrations in Neff reflects broader genome-wide patterns in translocation, indicating that integrated viral DNAs are subject to broader genomic forces shaping the *Acanthamoeba* genome. Across eukaryotes, sub-telomeric regions have been linked to relaxed selection, lateral gene transfer and transposable elements^53,54,55^, making them plausible landing pads for foreign DNA.

Ironically, the same processes that neutralize viral DNA over extended periods of time provide an opportunity for viral genes to very occasionally become functional. The translocation of DNA throughout the genome would sometimes allow viral genes to acquire appropriate transcription regulatory sequences and become expressed, and their integration in specific alleles may allow the healthy, original allele(s) to compensate for potential deleterious effects, providing a window of time for neofunctionalization. But successful acquisitions are overwhelmingly exceptions, judging from current patterns of virus-to-eukaryote transfers^12^.

### Broader implications across the tree of eukaryotes

How applicable is this three-step model across the eukaryotic tree of life? One major obstacle to generalization is the difficulty of comparing different genomic assemblies. Even between *A. castellanii* strains Neff and C3 we find striking divergences. Viral insertions in the C3 genome are smaller, less numerous, and less clearly biased towards sub-telomeric regions as in Neff. The extent to which these differences are an artefact of assembly problems linked to imperfect long-read sequence data and aneuploidy remains to be seen. We have shown that viral insertions in *Acanthamoeba* can be biologically complex and consequentially vulnerable to mis-assembly, and that viral sequences in C3 are disproportionately found on small scaffolds.

Some clear general patterns can nevertheless be observed across genomic data. Complete, partial and highly degraded viral genomes have all been found integrated in the genomes of diverse protists, indicating that there exists a spectrum of viral integration – from mirusvirus elements in *Aurantiochytrium* and other thraustochytrids^16^ to the ‘giant endogenous viral elements’ of the green *alga Chlamydomonas*^17^ and brown alga *Ectocarpus*^56^, to the scattershot integration described here in *Acanthamoeba*. Hypermethylation of fragmented viral regions has also been observed in the unicellular opisthokont *Amoebidium*^22^ which also shows evidence for recurrent, transient viral integration. While the model we outline is by no means the only existing pathway to integration, evidence suggests that it is likely very common. Understanding the interplay between methylation and integration will be particularly important to tease out variations in this model, thereby revealing differences in the way that epigenetics is likely to be utilized as a ‘tool’ by eukaryotic hosts to regulate the expression of foreign DNA in particular and resident genes in general.

We have shown that viral DNA insertions in *Acanthamoeba castellanii* genomes are numerous but transient, and do not contribute to the kind of genome bloat seen in many complex multicellular organisms or in dinoflagellates. The value, or lack thereof, that selfish DNAs such as those coming from viruses brings to their host is a long-standing debate. As the study of viral integration in protists comes to maturity, it becomes possible to recast debates around selfish DNA in a truly phylogenetically and comparatively rigorous context. Cells everywhere are constantly exposed to a barrage of foreign DNA. How they respond to it, and what they do with it, is a fundamental question for our understanding of evolution.

## Conclusion

We analyzed chromosome scale assemblies for the *Acanthamoeba castellanii* strains Neff and C3 and identified hundreds of new potentially viral sequences. Our analysis shows that viral insertions are largely transient and not conserved between the two genomes. Most conserved viral candidates are in fact likely eukaryotic genes which happen to have viral homologues due to eukaryote-to-virus lateral LGT. Viral insertions correspond to partial chunks of larger viruses and are highly methylated, with the largest two viral regions in the Neff genome apparently localized to a single heterochromatic structure. Multiple partial copies of viral insertions can be found throughout the genome, which may result from multiple insertions by the same virus or duplications following insertion. Although they are not mobile, the position of viral DNA fragments in the genome is likely dynamic. This dynamism is multifaceted and includes duplication of subsets of viral regions, chimeric regions combining multiple viral insertions, host-gene colonization of viral regions, and allele-specific variants of viral integrations. We synthesize these observations into a cohesive integration, suppression, and degradation model of the viral footprint in *Acanthamoeba*, which can be applied to and tested in other lineages. Integration may result from accidental recombination between the viral and host genomes during viral replication in the nucleus, and/or from complete endogenization of viral genomes into the host. Suppression further explains how the cell survives the initial infection and can retain the virus over multiple generations. Degradation permanently inactivates the viral DNA, shattering it into smaller pieces which, although they do not encode machinery for their own mobility, can be dispersed throughout the genome by mutational processes intrinsic to the host. Observed viral integrations represent transitory stages between insertion and deletion. Viral DNA in *Acanthamoeba* is thus likely the evolutionary by-product of recurrent close contact with viruses and does not appear to have long term effects on genome content, although it may be a facilitator of intra- and inter-chromosomal recombination events. Why some genomes appear to retain the large amounts of ‘junk’ they receive while others, like *Acanthamoeba*, delete them relatively quickly over evolutionary timescales, is a question that still needs to be addressed.

## Acknowledgements

We thank Cyril Matthey-Doret for providing raw nanopore sequence reads for the C3 strain of *Acanthamoeba*. Research in the Archibald Lab was supported by the Gordon and Betty Moore Foundation (GBMF5782) and a Discovery Grant (RGPIN-2019-05058) from the Natural Sciences and Engineering Research Council of Canada (NSERC). C.B. is supported by graduate scholarships from NSERC, the Nova Scotia Graduate Scholarship Program and Dalhousie University. M.J.C. was supported by graduate student scholarships from NSERC and Dalhousie University. L.S. and A.d.M. are supported by a European Research Council Starting Grant (ERC-StG 950230).

